# High-density, targeted monitoring of tyrosine phosphorylation reveals activated signaling networks in human tumors

**DOI:** 10.1101/2020.06.01.127787

**Authors:** Lauren E. Stopfer, Cameron T. Flower, Aaron S. Gajadhar, Bhavin Patel, Sebastien Gallien, Daniel Lopez-Ferrer, Forest M. White

**Affiliations:** Koch Institute for Integrative Cancer Research, Massachusetts Institute of Technology, Cambridge, MA; Center for Precision Cancer Medicine, Massachusetts Institute of Technology, Cambridge, MA; Department of Biological Engineering, Massachusetts Institute of Technology, Cambridge, MA; Program in Computational and Systems Biology, Massachusetts Institute of Technology, Cambridge, MA; Thermo Fisher Scientific, San Jose, CA; Thermo Fisher Scientific, Rockford, IL; Thermo Fisher Scientific, Precision Medicine Science Center, Cambridge, MA

## Abstract

Tyrosine phosphorylation (pTyr) plays a pivotal role in signal transduction and is commonly dysregulated in cancer. As a result, profiling tumor pTyr levels may reveal therapeutic insights critical to combating disease. Existing discovery and targeted mass spectrometry-based methods used to monitor pTyr networks involve a tradeoff between broad coverage of the pTyr network, reproducibility in target identification across analyses, and accurate quantification. To address these limitations, we developed a targeted approach, termed “SureQuant pTyr,” coupling low input pTyr enrichment with a panel of isotopically labeled, tyrosine phosphorylated internal standard (IS) peptides. Using internal standard guided acquisition, the real-time detection of IS peptides during the analysis initiates the sensitive and selective quantitation of endogenous pTyr targets. This framework allows for reliable quantification of several hundred commonly dysregulated pTyr targets with high quantitative accuracy, enhances target detection success rates, and improves the robustness and usability of targeted acquisition. We establish the clinical applicability of SureQuant pTyr by profiling pTyr signaling levels in human colorectal tumors using minimal sample input, characterizing patient specific oncogenic driving mechanisms. While in some cases pTyr profiles align with previously reported proteomic, genomic, and transcriptomic molecular characterizations, we highlight instances of new insights gained using pTyr characterization and emphasize the complementary nature of pTyr measurements with traditional biomarkers for improving patient stratification and identifying therapeutic targets. The turn-key nature of this approach opens the door to rapid and reproducible pTyr profiling in research and clinical settings alike and enable pTyr-based measurements for applications in precision medicine.

**Summary:** A targeted, mass spectrometry-based method, termed “SureQuant pTyr,” enables highly sensitive and reproducible profiling of tyrosine phosphorylation levels in human colorectal tumors and reveals dysregulated signaling networks for enhanced tumor characterization and biomarker identification.

## Introduction

Protein posttranslational modifications (PTMs) provide a fundamental mechanism to regulate protein function. The most common PTM, phosphorylation, is reversibly mediated by a network of protein kinases and phosphatases. Phosphorylation can cause conformation changes that activate or inactivate proteins, while also recruiting adaptor proteins and substrates that initiate downstream signaling cascades, thus altering the cell state (*1–3*). While over 250,000 unique phosphorylation sites have been reported, nearly all phosphorylation sites occur on serine and threonine residues, and less than ∼1% occur on tyrosine residues (*4–6*). Thus, deep profiling of tyrosine phosphorylation (pTyr)-mediated signaling requires pTyr enrichment and substantially higher sensitivity than standard phosphoproteomic or protein expression profiling approaches. Despite the rarity of pTyr, tyrosine kinases play a critical role in the signal transduction of pathways controlling proliferation, apoptosis, and survival, and their dysregulation through mutation, hyperactivation, or overexpression can lead to tumorigenesis (*7, 8*).

Many cancer therapeutics target oncogenic tyrosine kinases (*9*). While kinase inhibitors have demonstrated clinical success, identifying patients that may benefit from specific therapies remains challenging, as a majority of clinical molecular characterization efforts rely on genomic-based methods, which do not necessarily reflect protein or pathway activation status and are unable to capture the complex dynamics of innate and acquired therapeutic resistance (*10, 11*). Tyrosine phosphoprotein measurements have proven valuable in identifying aberrantly activated signaling pathways and characterizing therapeutic resistance mechanisms (*12–15*), which should provide biomarkers to help inform personalized therapies. Unfortunately, measuring low abundance tyrosine phosphorylated peptides remains challenging, particularly from limited amounts of sample material.

Existing methods to profile pTyr levels are well documented, but each requires a compromise between sensitivity, reproducibility, broad coverage, and quantitative accuracy. Phosphorylation site-specific antibodies have been applied in a variety of formats, including multiplex immunoassays and reverse phase protein arrays, among others. While these assays are relatively straightforward and reproducible, it remains difficult to measure low abundance targets and distinguish between similar phospho-epitopes on distinct proteins due to poor antibody specificity (*16, 17*). High sensitivity, mass spectrometry (MS)-based pTyr methods provide an attractive alternative, although each of the three typical data acquisition strategies has limitations. Data-dependent acquisition (DDA) or “shotgun” MS-methods offer deep sequencing of the tyrosine phosphoproteome without requiring previous knowledge of peptide targets, enabling novel discovery (*12, 18, 19*). However, DDA methods also result in inconsistent reproducibility of detected peptides, arising from stochastic sampling of precursor ions, and can be biased towards peptides of higher abundance (*20, 21*). Targeted methods like parallel or multiple-reaction monitoring (PRM/MRM) are well suited to quantify a known panel of peptides with high accuracy and reproducibility, but such traditional targeted acquisition schemes often require a tradeoff between the number of peptides that can be reliably measured and the sensitivity and selectivity of those measurements, restricting depth of coverage (*22*). These methods also commonly require complex method acquisition structures and peptide retention-time scheduling, which limits ease of use (*21*). Finally, pTyr data-independent acquisition (DIA) methods aim to improve run-to-run overlap while maintaining depth of coverage (*5, 23*). However the complexity of DIA spectra make quantitative accuracy challenging, and DIA methods have demonstrated lower sensitivity than PRM approaches, a critical consideration with low abundance, tyrosine phosphorylated peptides (*24*).

To address these limitations in existing pTyr profiling strategies, we describe a novel, high-density, targeted MS approach, termed “SureQuant pTyr,” that leverages isotopically labeled, tyrosine phosphorylated internal standard (IS) trigger peptides to efficiently guide MS acquisition in real-time. Adapted from traditional IS-PRM (*25*), the use of trigger peptides eliminates the need for retention time scheduling to expand the capacity of targetable nodes, allowing for the reliable and accurate quantification of several hundred tyrosine phosphorylated peptide targets commonly dysregulated in cancer. This platform accommodates low sample input for pTyr enrichment and utilizes commercially available pTyr enrichment reagents, nano-HPLC columns, and data acquisition method templates for a streamlined, “plug and play” implementation.

We apply this approach to profile the pTyr signatures of human colorectal cancer (CRC) tumor specimens to identify dysregulated signaling pathways and reveal potential drug targets not identified with genomic or other proteomic measurements, such as tumors susceptible to anti-epidermal growth factor receptor (EGFR) therapy. Furthermore, we demonstrate the tumor-extrinsic nature of pTyr profiling on tumor specimens, quantifying T cell activation levels on low abundance immune cell-specific pTyr sites, which may be an effective indicator of immune cell infiltration and immunotherapy response. With the reproducibility and sensitivity of SureQuant pTyr, we highlight the potential of this approach to be used in clinical settings to rapidly profile pTyr signaling as a complementary strategy to enhance biomarker identification and tumor characterization for applications in precision medicine.

## Results

### Targeted pTyr proteomic workflow utilizing internal standard driven data acquisition

In order to profile pTyr signaling events in cancer, we selected 340 tyrosine phosphorylated peptides to target and synthesized the corresponding synthetic isotope labeled (SIL) phosphopeptides to serve as ISs (Fig. 1A, data file S1). Selected peptides were primarily chosen from discovery analyses performed on a cohort of CRC samples; this list was then supplemented with additional peptides from pathways of interest. These sites include EGFR and T cell signaling pathway peptides, as EGFR inhibitors and immune checkpoint blockade (ICB) are two common therapies within CRC and other cancer types (*26, 27*). Selected pTyr sites primarily cover two branches of the kinome: tyrosine kinases and CMGC kinases (cyclin-dependent kinases (CDK), mitogen-activated protein (MAP) kinases, glycogen synthase kinases, and CDK-like kinases), which include tyrosine kinase signaling pathways relevant in cancer (*7*).

**Fig. 1.**
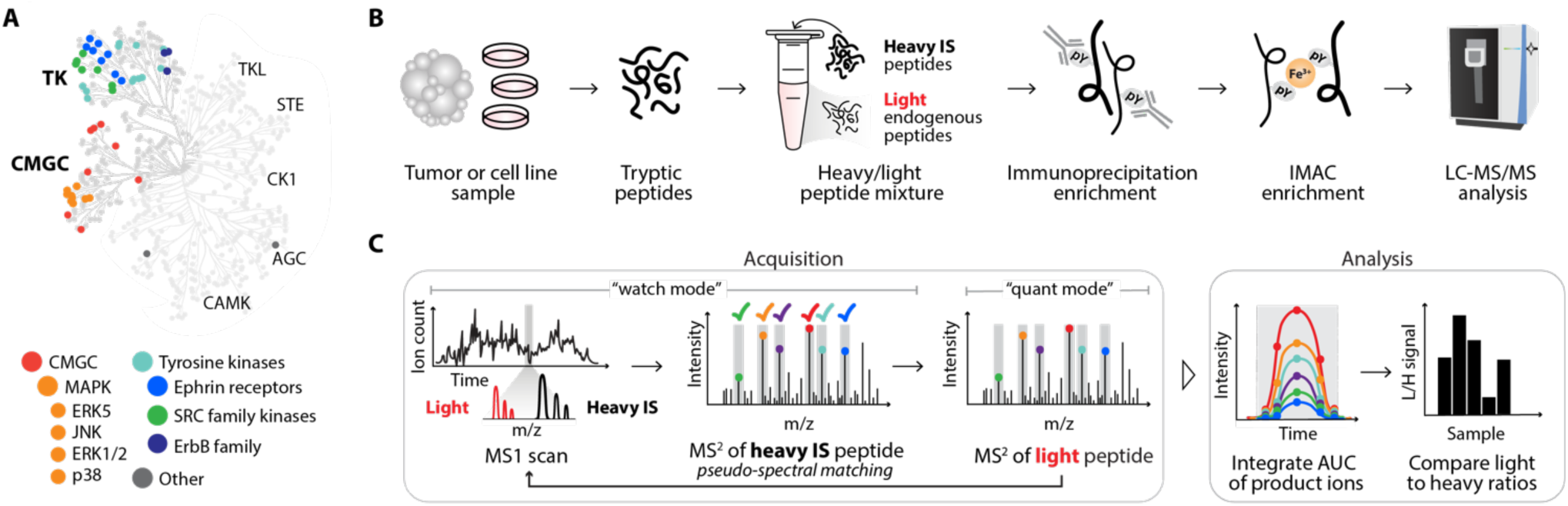
Platform for targeted pTyr analysis with IS-guided acquisition. (A) Human kinome tree with peptides selected for SureQuant pTyr analysis colored according to kinase group. (B) Sample processing workflow for pTyr enrichment and analysis. (C) Mass spectrometry acquisition method and analysis workflow for SureQuant pTyr IS-triggered quantitation.

For SureQuant pTyr analysis, tumors or cell line samples were first digested into tryptic peptides, and stable isotope-labeled, tyrosine phosphorylated IS (*i.e.*, “heavy”) peptides were added to the endogenous (*i.e.*, “light”) peptide mixture. (Fig. 1B). Both light and heavy pTyr peptides were subsequently isolated using two-step enrichment, with an immunoprecipitation against pTyr residues, followed by immobilized metal affinity chromatography (IMAC). Enriched light and heavy pTyr peptides were next analyzed by LC-MS/MS using a custom IS-triggered targeted quantitation method, leveraging the “SureQuant” acquisition mode native to the Orbitrap Exploris 480 MS (Thermo Scientific).

During SureQuant acquisition, the MS alternates between a “watch” mode and a “quantitative” mode (Fig. 1C). In watch mode, the MS continuously monitors for the presence of any heavy IS peptide. If an IS precursor ion is detected above a specified intensity threshold, a fast, low resolution MS^2^ scan is performed and pseudo-spectral matching against six pre-selected product ions is applied to verify the presence of the IS for enhanced selectivity. If the MS^2^ spectrum is a positive match, the MS initiates quantitative mode, triggering a high-quality MS^2^ scan of the light, endogenous peptide. With this framework, IS-guided acquisition ensures high selectivity, high sensitivity measurements of the endogenous peptide for enhanced data quality and reproducibility.

Product ions for both the heavy IS and light target peptides are monitored throughout the peptides’ chromatographic elution, and signal intensity is quantified by integrating the area under the curve for both light and heavy peptide product ions. Next, the ratio of light signal to heavy signal (L:H) is calculated, and L:H ratios are used for quantitative comparisons across samples. Adding IS peptides at defined concentrations prior to pTyr enrichment provides a number of additional benefits, serving as an embedded standard for concentration or copy number estimation using one-point calibration and enabling normalization across a theoretically unlimited number of samples and data-collection sites. IS peptides also double as a limit-of-detection control, as identification of the heavy IS but not the light peptide suggests the endogenous peptide was absent or below the limit-of-detection. Importantly, all parameters necessary to implement this workflow are readily determined in a single survey run analysis and can be used for all subsequent SureQuant analyses of the same peptide panel, streamlining assay implementation.

### SureQuant pTyr acquisition yields reproducible quantitation across replicate analyses

We first applied this workflow to measure pTyr levels in A549 lung carcinoma cells stimulated with epidermal growth factor as an in vitro control. We isolated light and heavy pTyr peptides from three technical replicate samples while varying the length of immunoprecipitation to assess the quantitative reproducibility of the SureQuant pTyr approach across replicate samples (Fig. 2A). Using a catenin delta-1 (CTTND1-pY904) peptide as an example, multiple MS^2^ scans were captured across the chromatographic peptide elution profiles for both the light and heavy peptide (Fig. 2B-C). Between replicates, the product ion intensities varied, with replicate 3 having over 3-fold higher signal intensity than replicates 1 and 2, likely due to a longer incubation time during immunoprecipitation (Fig. 2D). Despite this dissimilarity in intensities, the L:H ratios across replicates remained consistent (Fig. 2E), demonstrating the ability of this workflow to account for variation in sample handling and absolute intensities. In fact, across all quantified peptides the correlation coefficient (r^2^) between analyses was greater than 0.95 (Fig. 2F).

**Fig. 2.**
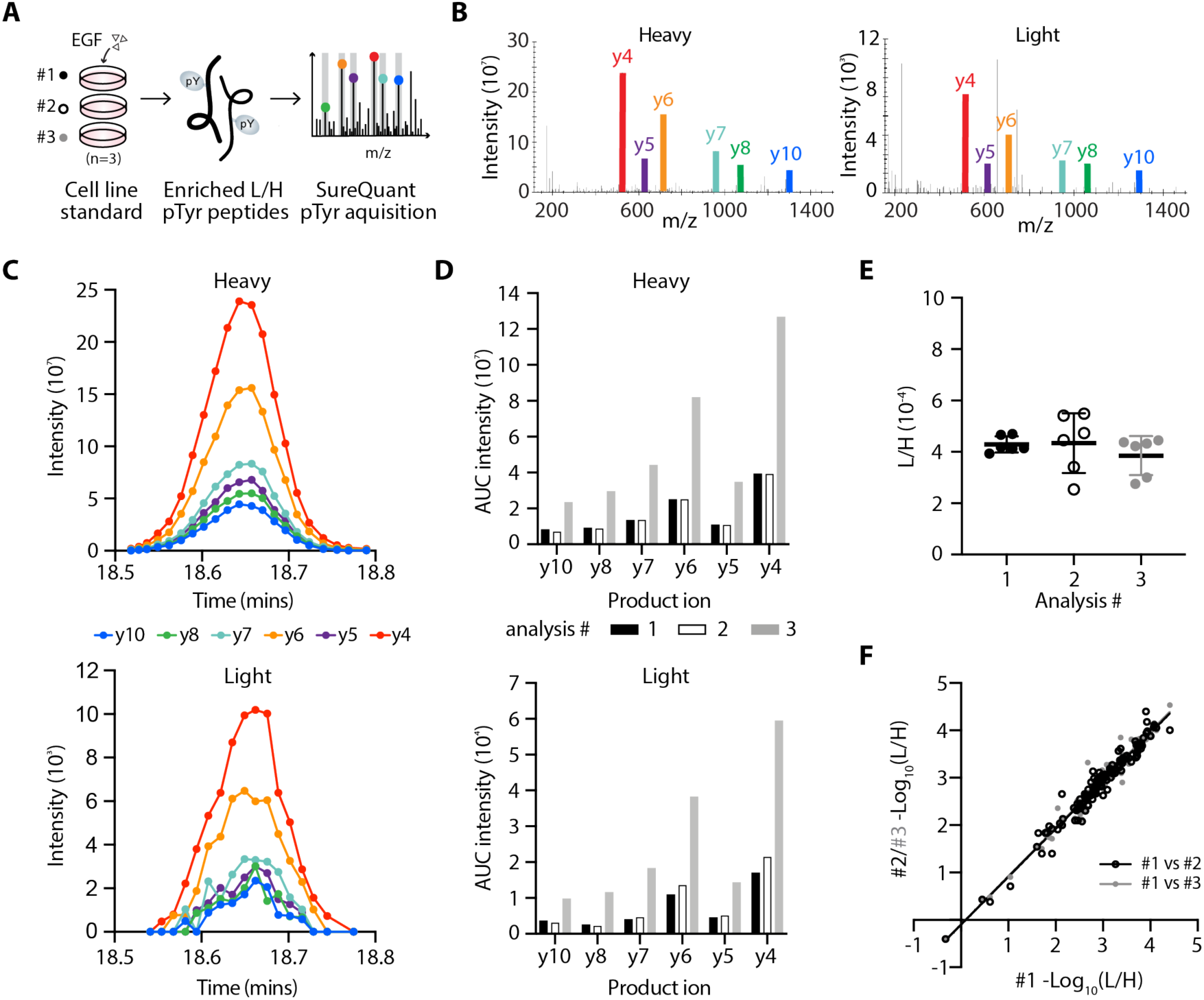
High quantitative reproducibility is achieved with SureQuant pTyr. (**A**) Experimental setup. IS-trigger peptides were added to three biological replicates of A549 cell lysate stimulated with epidermal growth factor (EGF). Enriched light (L) and heavy (H) pTyr peptides were analyzing using SureQuant pTyr acquisition. (**B**) MS/MS spectra from analysis #3 of the heavy (left) and light (right) CTTND1 peptide, SLDNN[pY]STPNER-pY904, at peak intensity, where [pY] denotes the residue position with pTyr modification. Monitored product ions are uniquely colored and labeled with b/y ion. (**C**) Ion intensity over time for the 6 heavy (upper) and light (lower) product ions from CTTND1 pY904 in analysis #3. Each MS/MS event is represented by a point. (**D**) Integrated area under the curve (AUC) intensities for each product ion in C. Bar color corresponds to analysis #. (**E**) Ratios of light to heavy signal intensity (L/H) of CTTND1 pY904 for each analysis, where each point represents the L/H value of a single product ion. (**F**) Correlation of (L/H) signals across 127 peptides between analysis #1 and #2 (r^2^=0.96, black) and analysis #1 and #3 (r^2^=0.97, grey).

### Human colorectal tumors show distinct pTyr signatures

Thirty-one human CRC tumors that were previously characterized in a proteogenomic analysis by Vasaikar et al. (*15*) were selected for SureQuant pTyr profiling. The previous study included a global phosphorylation analysis, but only 16/2183 sites (0.07%) measured across all 31 tumors were tyrosine phosphorylated, underscoring the need for pTyr-specific enrichment. Using our panel of 340 tyrosine phosphorylated ISs, we collectively detected 336 heavy peptides, representing 99% of the assay panel, and 325 light peptides, representing 96% of endogenous peptides from the assay panel across the tumor cohort (Fig. 3A, fig. S1A). The four unmeasured heavy peptides exhibited fluctuating signal from run to run and did not systematically reach the signal intensity threshold defined in the initial survey analysis, while the eleven unquantifiable light peptides are assumed to be below the limit-of-detection, as the corresponding heavy peptide was detected. Across all tumors, an average of 91% of heavy peptides and 78% of light peptides were identified, highlighting the reproducibility of the method. While we did not see complete coverage of our panel in every tumor, this result was expected as pTyr peptides are often present at low levels, and some of the peptides included in the panel were hypothesis driven. For example, we included a T cell signaling peptide from ZAP70 (pY292) which was not identified in the discovery analyses but was quantifiable in 16/31 tumors. Due to the tumor-specificity of some of the signaling nodes, we analyzed the pTyr signaling in two ways. First, L:H ratios of peptides identified across all 31 tumors were quantified and z-score normalized (“full matrix*”)*. Second, the L:H ratios for all sites identified in at least 50% of tumors were z-score normalized to expand the datasets for individual tumor signaling analyses (“expanded matrix”).

**Fig. 3.**
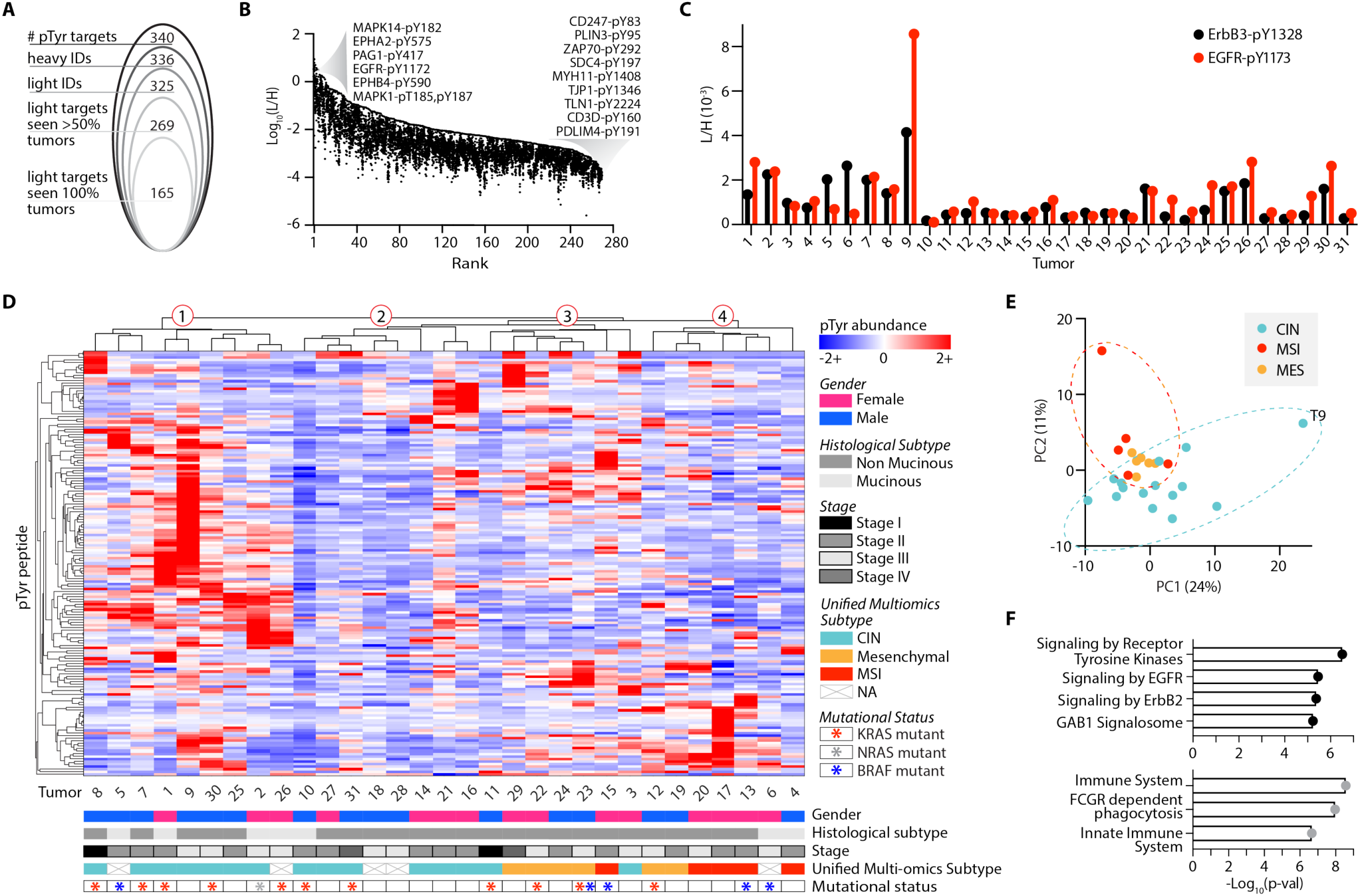
Targeted pTyr analysis highlights CRC tumor heterogeneity. (**A**) Peptides identified and quantified across 31 tumors. (**B**) Distribution of tumor L:H ratios with peptides rank ordered from highest to lowest maximum abundance. Annotated peptides labeled by source protein and residue position with pTyr modification have the maximum and minimum abundances. (**C**) Light to heavy signal intensity ratios (L/H) for ErbB3 peptide (black), SLEATDSAFDNPD[pY]WHSR, and EGFR peptide (red), GSTAENAE[pY]LR, where [pY] denotes the residue position with pTyr modification. (**D**) Peptide and tumor hierarchical clustering (distance metric = correlation), where pTyr abundance values are z-score normalized light to heavy signal ratios. (**E**) Tumors plotted by principal component 1 (PC1) and PC2 score, colored according to unified multi-omics subtype. (**F**) Significantly enriched reactome pathways from the top 20 peptides derived from unique proteins on PC1 (top, black) and PC2 (bottom, grey). Significance values are FDR adjusted.

The peptide L:H ratios spanned six orders of magnitude, with just 0.7% of sites having a L:H ratio above 1, indicating that most of the endogenous pTyr peptides fell below the nominal amount (∼1 pmol) of IS used, while a few exceeded this amount (Fig. 3B). These highly abundant sites include MAPK3/1 (ERK1/2), EGFR, and MAPK14 (p38*α*), each of which are implicated in oncogenesis (*7, 9*). Sites with the lowest L:H ratios include several T cell signaling associated peptides (CD3ζ, CD3δ, ZAP70), consistent with our hypothesis that a minority of cells in these tumors are infiltrating immune cells.

The variation biological variation in peptide pTyr levels between tumors is evident upon comparing the L:H ratios measured by this platform. For example, EGFR and ErbB3, two receptor tyrosine kinases (RTKs) in the epidermal growth factor receptor family, appear to have coordinated levels of receptor phosphorylation in some tumors (Tumor 2 (T2), T7, T8, T21, T25), while others have differential levels (Fig. 3C). ErbB3 is a non-autonomous receptor, requiring dimerization with another ErbB family member or RTK for phosphorylation. Thus, the higher ErbB3 and lower EGFR phosphorylation levels of T5 and T6 suggest ErbB3 may be dimerizing with another ErbB family member or activated RTK. Alternatively, T1, T26, and T30 show the opposite trend, implying ErbB3 is playing a less dominant role in driving ErbB family signaling in these tumors. To assess whether observed pTyr abundance differences could be explained by variation in the overall amount of pTyr signal among tumors, potentially due to differences in protein loading or sample processing, we evaluated the distribution of L:H ratios across tumors (fig. S1B). Only two tumors, T9 and T10 had a significantly higher and lower distribution, respectively, of L:H ratios from the mean signal across tumors, suggesting differences in pTyr levels are indicative of biological variation as opposed to experimental variation.

To visualize the pTyr signaling profiles across tumors, phosphorylation sites quantified in all tumors were analyzed by hierarchical clustering (Fig 3D). Two clear findings emerge from this analysis: each tumor possesses a unique pTyr signature, and tumor clustering is not readily explained by phenotypic information such as gender, histological subtype, or tumor stage (data file S2). Previous work by Vasaikar et al. assigned each tumor in our panel to one of three unified multi-omics subtypes (UMS), characterizing tumors with microsatellite instability and hypermutation (“MSI”), chromosomal instability (“CIN”), and evidence of epithelial-to-mesenchymal transition (“mesenchymal”), based off of previous proteomic, genomic, and transcriptomic-based classifications developed for CRC tumors (*15, 28, 29*). These classifications revealed some stratification with hierarchical clustering: CIN tumors are primarily located in clusters one and two, whereas a majority of mesenchymal and MSI tumors group together in clusters three and four, respectively. Still, hierarchical clustering of tumors with the same UMS illustrates the high degree of individuality in each tumor’s pTyr signature, even within co-clustering subtypes (fig. S1C).

To understand which pTyr sites drive the UMS clustering of tumors, we utilized principal component analysis (PCA) (Fig. 3E). Principal component 1 (PC1), explaining 24% of the total variance, primarily separates T9 from the remaining tumors and is driven by T9’s high pTyr levels of EGFR signaling peptides (Fig. 3F, fig. S1D). Interestingly, PC2, explaining 11% of the total variance, separates CIN tumors from the MSI and mesenchymal tumors. The 20 highest scoring pTyr peptides derived from unique proteins on PC2 show enrichment for pathways related to innate immunity (Fig. 3F, fig. S1E). Vasaikar et al. found that MSI and mesenchymal tumors had higher levels of immune cell infiltration, in agreement with our pTyr findings.

### Tumor-specific pathway analysis reveals enriched signaling pathways

We next performed a correlation analysis on the full peptide matrix and clustered peptides on this basis to identify groups of co-regulated peptides across tumors (Fig. 4A). A protein-protein interaction network analysis on selected clusters revealed significantly enriched pathways and processes (fig. S2A). These included pathways related to immunity in cluster 1 (Fig. 4B), as well as cytoskeletal and actin binding proteins in cluster 2 (Fig. 4C). Cluster 3 maps to ErbB and Ras signaling pathways, (Fig. 4D), along with migration signaling pathways like adherins junctions, focal adhesions, and RAP1 signaling (Fig. 4E). Using these findings, we curated a custom library of twelve gene sets and performed a tumor-specific pathway enrichment analysis (TPEA). Phosphorylation site source proteins from the expanded values matrix were rank ordered and used to identify tumors with positive or negative enrichment in the selected pathways and biological processes relative to the other tumors (Fig. 4F), showcasing the pathway level information obtained with SureQuant pTyr. For example, T16 and T10 have significant positive and negative enrichment in actin binding phosphopeptides, respectively, and correspondingly have the highest and lowest phosphorylation levels of peptides identified in cluster 2 (fig. S2B). While some findings were redundant with insights obtained with hierarchical clustering, TPEA also identified signaling level similarities between tumors that were not obvious with clustering. For instance, T13 and T29 both have significant positive enrichment of RAP1 signaling but clustered separately in Fig. 3D.

**Fig. 4.**
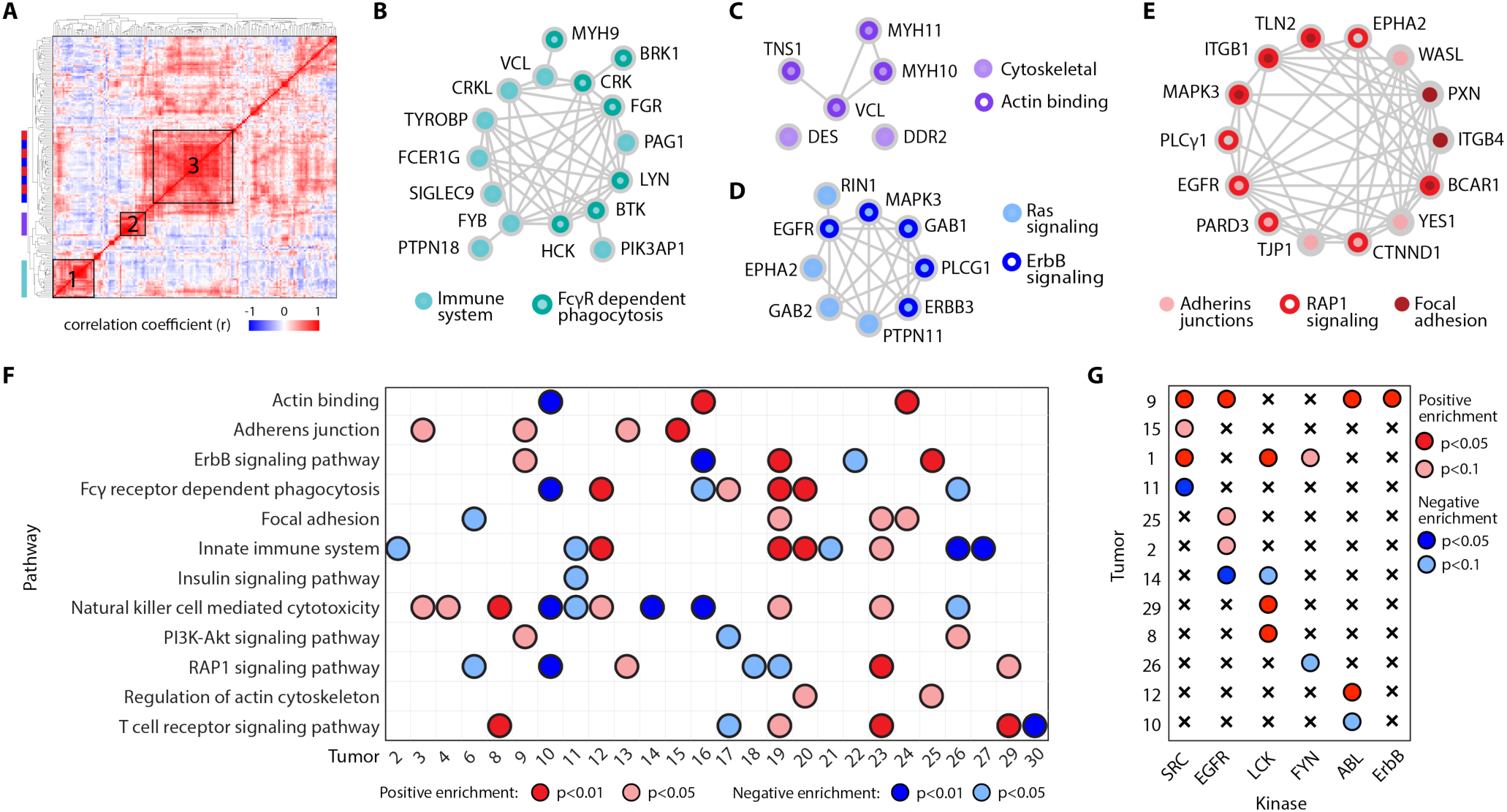
Differential pTyr levels identify tumors with significant pathway enrichment. (A) Hierarchical clustering based on the correlation coefficients between phosphosites across all tumors (distance metric = correlation). (B-E) Protein-protein interaction network of peptides within cluster 1 (B), cluster 2 (C) and cluster 3 (D-E). Node color(s) maps peptides to enriched pathway(s). (F) Significantly enriched pathways among tumors using tumor-specific pathway enrichment analysis. (G) Significantly enriched kinase-substrate interactions within tumors. Significance (p-value) and directionality indicated by color, q < 0.25 for all enrichment analyses. Tumors that did not have any significant enrichment are not shown.

Additionally, we applied kinase-substrate enrichment analysis (KSEA) to each tumor which, in contrast to TPEA, uses site-specific information to identify the enrichment of phosphorylated kinase substrates to infer kinase activity (Fig. 4G) (*30*). The results were complementary in some cases, with T9 showing an enrichment in ErbB signaling pathways with TPEA and ErbB substrates with KSEA, but KSEA also revealed novel findings. T1 did not contain any significantly enriched pathways, but showed significant enrichment in SRC family kinase substrates including SRC, LCK, and FYN, which have been explored as therapeutic targets in metastatic CRC (*31*). To better understand each tumor’s unique pTyr profile and identify therapeutically targetable nodes, we next examined the sites driving pathway enrichment, focusing first on ErbB signaling and EGFR phosphorylation status.

### ErbB phosphorylation levels identify candidates for anti-EGFR therapy

EGFR is expressed in a majority of CRC, and its overexpression in many cancer types has been tied to more aggressive phenotypes and poor clinical prognosis, highlighting EGFR inhibitors as a promising therapeutic target (*32*). Indeed, several anti-EGFR agents have been approved for CRC clinical use, though treatment is currently only recommended for patients with wild type KRAS/NRAS/BRAF, as mutations in these genes have been shown to confer EGFR inhibitor resistance (*33–35*). Disappointingly, anti-EGFR agents are only effective in a fraction of qualifying patients, and those that do respond often still develop therapeutic resistance (*36*). As EGFR expression levels have not been shown to correlate with clinical response to EGFR inhibitors (*37*) and RAS mutational status remains the principle biomarker for EGFR inhibitor efficacy, we hypothesized that measuring pTyr levels on EGFR and ErbB family signaling pathways could provide a more direct readout of EGFR activation status, thereby improving identification of those who may benefit from EGFR inhibition.

We identified three tumors with significant positive enrichment of the ErbB signaling pathway (T19, T25, and T9), and two with significant negative enrichment (T16 and T22). T19 has low pTyr levels of the ErbB family receptors, with pathway enrichment instead driven by common downstream signaling nodes including ERK1/2 phosphorylation (fig. S2C), and therefore, T19 was excluded from subsequent analyses. Neither T25 nor T9 contained a RAS/RAF mutation, making both eligible for anti-EGFR therapy under existing biomarker criteria. T25 displayed high levels of EGFR phosphorylation, suggesting T25 may be a good candidate for an EGFR-inhibiting antibody like cetuximab (Fig. 5A). Alternatively, T9 had high levels of three ErbB RTKs: EGFR, ErbB2, and ErbB3, indicating EGFR inhibition alone may not be sufficient for T9, as ErbB2 amplification is predictive of anti-EGFR therapy resistance (*27, 37*). Instead, T9 may benefit from treatment with a pan-ErbB inhibitor like lapatinib, or combination therapy with cetuximab and the ErbB2 inhibitor, pertuzumab.

**Fig. 5.**
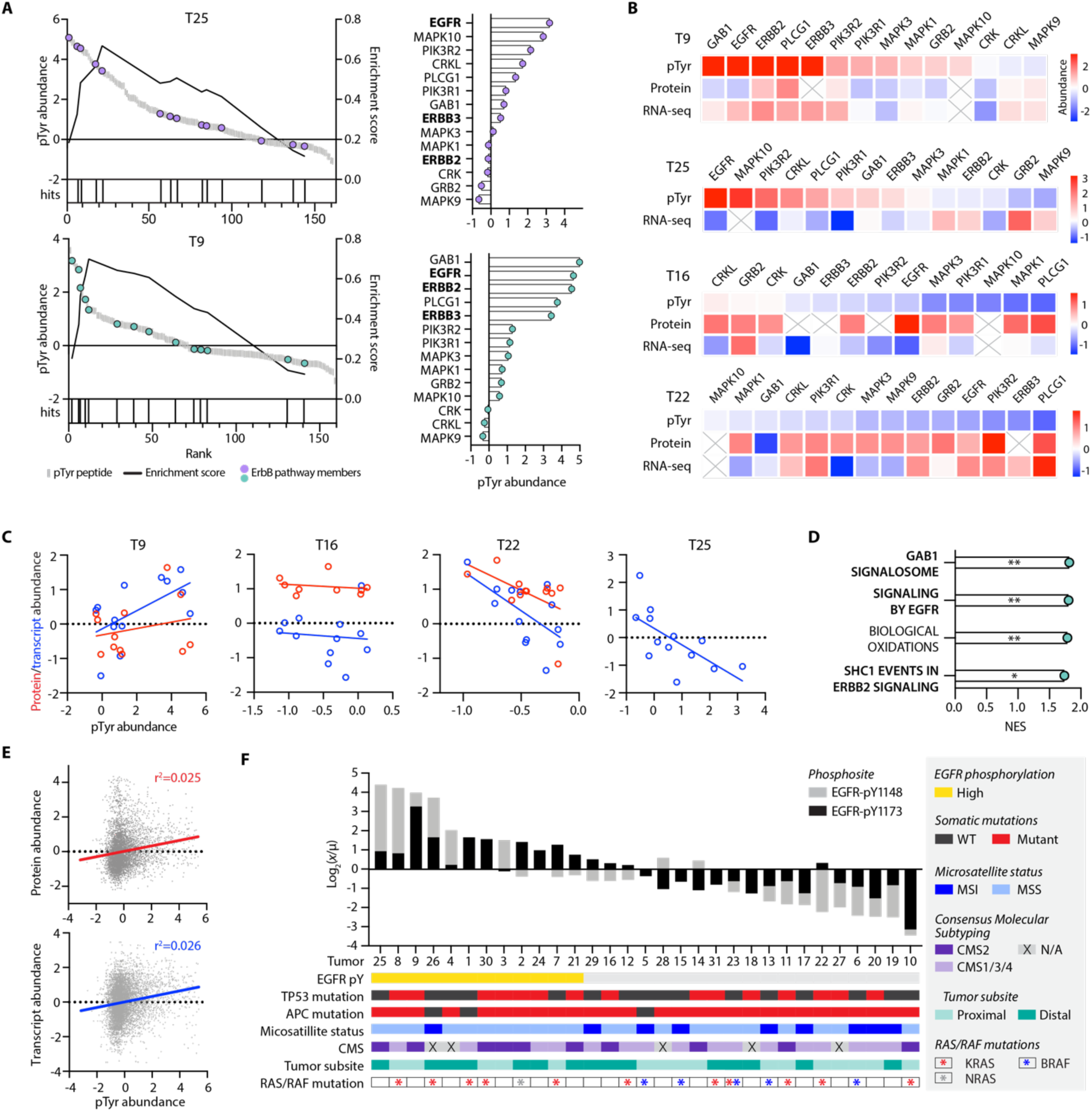
EGFR phosphorylation levels identify candidates for anti-EGFR therapy. (**A**) Enrichment plots (left) of ErbB signaling pathway in T25 (purple) and T9 (teal). Peptide rank (x-axis) versus pTyr abundance is plotted on the left y-axis, and the running enrichment score is plotted on the right y-axis. Each hit signifies a pTyr source protein present in the ErbB signaling pathway library. All pTyr peptides identified in the ErbB signaling pathway and their corresponding pTyr abundance (right), with ErbB family receptors annotated in bold. pTyr abundance values are z-score normalized light to heavy signal ratios. (**B**) Z-score normalized pTyr abundance, protein expression and transcript expression levels of ErbB signaling pathway members. (**C**) Correlation between pTyr abundance and protein (red) or transcript (blue) expression of ErbB signaling pathway members. Protein and transcript abundance values are z-score normalized. Correlation coefficients for pTyr vs. protein for T9, T16, and T22 are r^2^= 0.06, 0.05, 0.29, respectively. Correlation coefficients for pTyr vs. gene expression for T9, T16, T22, and T25 are r^2^= 0.35, 0.01, 0.38, and 0.43, respectively. Protein expression data for T25 was unavailable. (**D**) Normalized enrichment score (NES) for positively enriched reactome pathways in T9 using RNA-seq data. *= p<0.05, **= p<0.01, FDR q-value < 0.05 for all. (**E**) Correlation between all pTyr sites in the full matrix and corresponding protein expression (top) and gene expression (bottom) levels. All values are z-score normalized. (**F**) Cumulative pTyr signal, calculated as the ratio of tumor light to heavy pTyr signal (*x*) to the mean light to heavy signal (*μ*) across tumors, log2 transformed for two EGFR phosphopeptides rank ordered from highest to lowest signal. Tumor specific annotations are indicated by color, and pY1148 and pY1173 denote the EGFR residue position with pTyr modification.

We next sought to determine whether these findings were predictable based on available transcriptomics and proteomics data for these same tumors (*15*). Looking specifically at the sites driving ErbB enrichment, we observed a weak positive correlation between pTyr levels and corresponding gene expression levels in T9 but found no correlation with protein expression (Fig. 5B-C). T16 had no correlation between pTyr levels and gene/protein expression, whereas T25 and T22 surprisingly showed a weak negative correlation. In line with these findings, gene set enrichment analysis (GSEA) of RNA-seq data from T9 identified significant enrichment in EGFR signaling genes, along with downstream pathways of EGFR activation including SHC1 and GAB1 signaling (Fig. 5D, fig. S2D). However, GSEA from T25, T16, and T22 showed no significant enrichment in EGFR/ErbB related signaling pathways using protein or gene expression data. In fact, an analysis of all pTyr sites and their corresponding protein and gene expression levels yielded no correlation (Fig. 5E), which taken together demonstrates the difficulty of using transcript expression or protein expression data to infer pTyr signaling dynamics and pathway activation.

Identifying anti-EGFR therapy candidates using TPEA requires enrichment among multiple nodes within the ErbB signaling pathway to achieve significance. To identify anti-EGFR therapy candidates that may have been missed using TPEA, we focused on two EGFR peptides containing autophosphorylation sites, pY1148 and pY1173, which were most commonly quantified across tumors as an analogous approach. We identified twelve tumors with EGFR phosphorylation levels at least 1.5-fold higher than the mean in either or both pTyr sites, termed “EGFR-high” (Fig. 5F). Half of EGFR-high tumors have a RAS mutation rendering them ineligible for anti-EGFR therapy, but the remaining six wild-type EGFR-high tumors (T3, T4, T21, T24, and previously identified T9 and T25) may be appropriate candidates. Similar to earlier findings, GSEA of transcriptomic and proteomic datasets for T3, T4, T21, and T24 did not identify enrichment in EGFR signaling pathways, highlighting the novel insight provided by pTyr profiling.

Beyond RAS mutational status, several other genomic and phenotypic classifications have been correlated with response to EGFR inhibitors, including tumors with a mutation in both *TP53* and *APC*, microsatellite stable (MSS) status, distally located CRC tumors, and those classified as consensus molecular subtype 2 (CMS2), an additional CRC molecular classification system (*27, 29, 38*). Of the six wild-type EGFR-high tumors, only T21 matches these additional genomic criteria. In contrast, T18 and T16 possess all of the described biomarkers, but also have lower levels of EGFR phosphorylation, indicating an alternative therapy may be more efficacious.

These results suggest that pTyr analysis can provide critical information regarding target activation that, when combined with genomic characterization, may improve patient stratification for targeted therapeutics.

### T cell phosphorylation levels suggest tumor immune cell activation and infiltration status

Following the demonstrated success of ICB in other solid tumors, immunotherapy has emerged as another therapeutic avenue in CRC with several ICB therapies approved for clinical use (*39*–*41*). However, efficacy of ICB in CRC has been limited to mismatch-repair deficiency and microsatellite instability classified tumors (dMMR-MSI), which typically have higher immune cell infiltration and mutational burden than MMR proficient, microsatellite stable (pMMR-MSS) tumors, increasing their susceptibility to ICB therapy (*42, 43*). Nevertheless, dMMR-MSI tumors represent a minority (∼15%) of CRCs (*44*) and overall response rates in recent clinical trials ranged from 30-55%, emphasizing the need for additional biomarkers of ICB efficacy (*39–41*).

Unlike other cancers, PD-L1 expression is not predictive of ICB response in CRC (*39*). Still, better response rates have been observed in tumors with higher levels of CD8+ tumor-infiltrating lymphocytes, regardless of microsatellite status (*45*). With this in mind, we investigated whether we could identify patients with high CD8+ T cell infiltration using the pTyr levels of immune cell-specific peptides to estimate ICB responsiveness.

The SureQuant pTyr IS panel contained T cell signaling-specific peptides derived from the T cell receptor CD3δ/γ, along with the T cell co-receptor CD3ζ (CD247) and zeta-chain associated protein kinase (ZAP70). These sites and other downstream signaling nodes comprised the T cell signaling pathway gene set used for TPEA; using this gene set we identified six tumors with significant enrichment: four positive (T8, T19, T23, and T29) and two negative (T30 and T17) (Fig. 4F, fig. S4). Examining the phosphosites driving enrichment, we observed high levels of CD3ζ phosphorylation across multiple pTyr sites and increased ZAP70 phosphorylation in T8 and T29, relative to the other tumors (Fig. 6A). Both tumors also show elevated LCK phosphorylation, a SRC kinase which phosphorylates immunoreceptor tyrosine-based activation motifs (ITAMS) on TCR/CD3 substrates including CD3ζ and ZAP70. Similarly, KSEA identified significant positive enrichment of LCK substrate phosphorylation in T8 and T29 (Fig. 4G). In contrast, T cell signaling enrichment in T19 and T23 was primarily driven by downstream signaling nodes non-specific to T cells (p38 and ERK), which highlights the importance of evaluating pathway enrichment on a site-specific level.

**Fig. 6.**
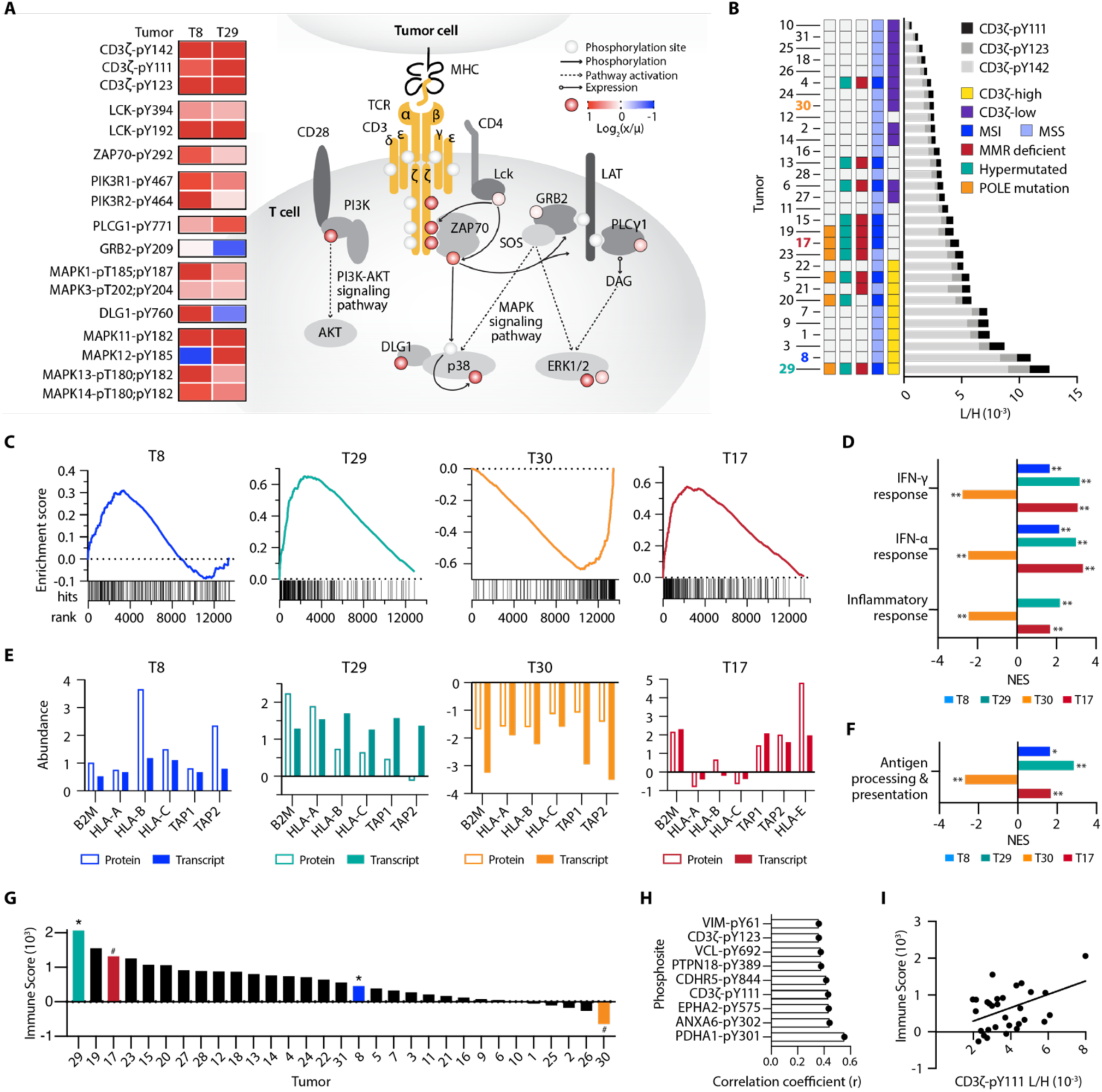
pTyr signatures of T cell signaling pathway peptides. (A) pTyr signal of T-cell signaling peptides, calculated as the ratio of tumor light to heavy pTyr signal (*x*) to the mean light to heavy signal (*μ*), for T8 and T9. Phosphorylation levels on the signaling diagram (colored circles) correspond to T8, and pY denotes the residue position with pTyr modification. (B) Cumulative CD3ζ pTyr levels (light to heavy signal) from three CD3ζ pTyr peptides. Tumor-specific biomarker statuses are indicated by color. (C) Gene set enrichment analysis (GSEA) plots for IFN-γ response, with gene rank (x-axis) versus running enrichment score (y-axis). Each hit signifies a gene present in the gene set. (D) Normalized enrichment scores (NES) from GSEA for selected significantly enriched pathways *p<0.05, **p<0.01, q<0.25 for all. (E) Z-scored normalized protein and transcript expression levels of antigen presentation genes. (F) GSEA NES for antigen processing and presentation gene ontology gene set (GO:0019882), *p<0.05, **p<0.01, q<0.25 for all. (G) Immune score of tumors from Vasaikar et al., * = positively enriched # = negatively enriched in pTyr T cell signaling peptides. (H) Unique pTyr peptides with significant positive correlation to immune score (p < 0.05). (I) Correlation between immune score and the L/H signal of CD3ζ-pY111, r^2^ = 0.19.

Consequently, we applied a parallel, site-specific framework as used in the phospho-EGFR analysis to evaluate three CD3ζ phosphosites as a marker for T cell infiltration. We identified ten tumors with at least a 1.5-fold increase in CD3ζ phosphorylation relative to the mean, similarly termed “CD3ζ-high,” and twelve tumors with at least 1.5-fold lower phosphorylation, “CD3ζ-low” (Fig. 6B). Only two CD3ζ-high tumors (T29 and T5) were classified as dMMR-MSI, while the others were pMMR-MSS. Mutations in DNA polymerase epsilon (POLE) have also been correlated with increase CD8+ T cell infiltration (*46*), but assessing POLE mutational status only identifies one additional tumor (T20) classified as CD3ζ-high. Other tumors including negatively enriched T cell signaling T17 and CD3ζ-low T4 may have been considered for ICB based on dMMR-MSI status, but low pTyr levels of CD3ζ suggest that they may not be strong candidates for ICB based on lack of activated T cell signaling within the tumor.

We examined whether transcript expression or protein expression datasets would similarly identify T cell signaling pathway enrichment among positively enriched T8/T29 and negatively enriched T30/T17, but this analysis yielded no significant findings. However, GSEA against the cancer hallmarks database using RNA-seq data identified interferon-γ (IFN-γ), IFN-α, and immune response pathways as significantly positively enriched in T8 and T29, and significantly negatively enriched in T30 (Fig. 6C and D). Tumor-infiltrating lymphocytes are the primary source of IFN-γ production in tumors and stimulates IFN response genes and immune activation, in agreement with our T cell signaling TPEA results (*47*). We next looked at expression levels of antigen presentation machinery, hypothesizing that expression levels would mirror the directionality of T cell signaling and IFN-γ enrichment. Supporting this notion, we see corresponding increased (T8, T29) or decreased (T30) gene and protein expression levels of class I major histocompatibility complex (MHC-I) genes (HLA-A/B/C, β2M) and both transporters associated with antigen processing (TAP1/2) subunits (Fig. 6E and F).

Intriguingly, negative T cell signaling enriched T17 has positive enrichment of IFN-γ/α, inflammatory response, and antigen processing and presentation genes, in contrast to the trend observed in the other tumors (Fig. 6C-F). Analysis of antigen presentation protein expression levels shows increased β2M and TAP1/2, but lower levels of classical HLA-A/B/C alleles with high expression of the non-classical allele, HLA-E. Similar to classical HLAs, HLA-E can be modulated by IFN-γ (*48*), but high expression of HLA-E can function as an inhibitory signal towards other immune cell types, attenuating tumor cell susceptibility to T cell mediated killing as a mechanism of immune escape (*49, 50*). Therefore, despite T17’s enrichment for IFN-γ/antigen presentation and biomarker status, a deeper analysis of the genomic and proteomic data aligns with our pTyr-based assessment of T17 being a poor candidate for ICB therapy.

Previous analyses by Vasaikar et al. provided their own estimate of immune infiltration by using a gene expression signature to assign each tumor an “immune score,” (Fig. 6G) representative of the fraction of immune cells within a tumor sample (*15, 51*). Tumors with the highest immune scores were largely dMMR-MSI and/or hypermutated, including negative T cell signaling enriched T17, while positively enriched T8 had an average score. To determine whether specific pTyr sites in our data were associated with tumor immune score, we assessed the correlation between these data. Although two CD3ζ pTyr sites were significantly correlated with the immune score, the correlations were weak, and other significantly correlated sites were not directly associated with T cell or immune function (Fig. 6H and I). Taken together, these data further highlight the additional information provided by directly measuring activation status with SureQuant pTyr and suggest pTyr measurements applied in conjunction with genomic and/or proteomic data may improve identification of potential ICB or other immunotherapy responders, beyond classical MMR/MSI biomarkers.

## Discussion

pTyr measurements are well suited to directly read out signaling network activation status and have the potential to identify therapeutically targetable protein kinases or signaling pathways in disease. To address the limitations of traditional shotgun and targeted MS-based pTyr approaches which require compromise between reproducibility, broad coverage, and quantitative accuracy, we developed SureQuant pTyr. This approach leverages isotopically labeled IS trigger peptides to guide the acquisition of endogenous pTyr peptides in real-time for enhanced sensitivity and selectivity. In addition, SureQuant pTyr addresses significant analytical challenges compared to other MS-based approaches, as it does not rely on retention time scheduling, thus maximizing the number of targetable peptides and reducing the complexity of assay development. We utilized a panel of 340 tyrosine phosphorylated IS peptides to obtain highly reproducible, high-density coverage of pTyr signaling pathways implicated in cancer, while achieving accurate quantitation.

We established the quantitative reproducibility of SureQuant pTyr, by performing targeted pTyr profiling on three replicate in vitro samples. While peptide intensity responses varied across analyses and thus may have confounded label-free analysis, L:H ratios were consistent. The high quantitative reproducibility achieved with the SureQuant IS-triggered targeted workflow has many benefits, including the ability to readily analyze signaling network dynamics under various conditions (*52*), or to compare the signaling state of a patient derived tissue over time as therapeutic resistance or metastases develop (*53*). Furthermore, using a set of reference standards for quantitation enables comparisons across research projects and data collection sites, paving the way for large scale, multi-site studies using pTyr levels for disease characterization.

To highlight the potential clinical utility of this platform, we applied SureQuant pTyr to measure pTyr signaling levels in CRC patient tumor tissues. Our analyses identified tumors with elevated EGFR and ErbB signaling levels, as well as tumors with high levels of innate and adaptive immune cell infiltration. While some of our results reinforced complementary genomics-based classifiers used for treatment selection, in other cases our pTyr signaling data revealed therapeutic opportunities in tumors that would have been missed by traditional biomarkers. These results demonstrate the power of pTyr characterization as a complementary approach for selecting treatment strategies. Furthermore, this platform only requires 800 μg of sample input material, less than a standard 14G needle biopsy, making it highly amenable to clinical sample profiling (*20*).

While our initial study utilized just 340 pTyr targets selected with CRC application in mind, the current method framework could be applied to an alternate or expanded panel of peptides for deeper profiling of the tyrosine phosphoproteome for applications in cancer research or other non-oncological settings where dysregulated kinase signaling also plays a role. In addition, coupling SureQuant acquisition with isobaric labeling would greatly increase assay throughput, allowing for up to 16 samples analyzed simultaneously (*54*) while further decreasing sample input material required (*53*).

Though this assay is not approved by the Clinical Laboratory Improvement Amendments (CLIA) for clinical use at this time, implementation of targeted MS in clinical settings is beginning to emerge (*55*). Importantly, while many targeted workflows require complex method structures and customized MS platforms, all aspects of pTyr SureQuant were performed with commercially available nano-HPLC columns, enrichment reagents, method templates, and instrumentation, thereby allowing for simplified implementation in other research or clinical settings. Executing this workflow simply requires the IS peptide mixture and a single survey analysis to determine intensity thresholds for IS peptide triggering, offering a turn-key solution for targeted pTyr profiling.

Improvements to assay accuracy, reproducibility, and ease of use may open new doors in the clinical setting to use utilize pTyr signatures in conjunction with existing technologies to obtain novel insights. Overall, we propose the broad application of targeted pTyr profiling with SureQuant pTyr in research and clinical settings can aid in improving patient stratification and biomarker characterization, identification of drug targets, and designing personalized therapies in the context of oncology and beyond.

## Materials and Methods

### Cell lines

Lung cancer cell line A549 (CCL-185) was purchased from ATCC and routinely tested for mycoplasma contamination (Lonza). Cells were cultured in RPMI-1640 (Gibco) supplemented with 10% FBS (Gibco), 1% penicillin/streptomycin (Gibco) and maintained at 37°C, 5% CO2. Prior to harvesting, cells were stimulated with 5 nM EGF (PeproTech) for 5 minutes.

### Tumor samples

Colon cancer tissue samples were obtained as tumor curls through the Clinical Proteomic Tumor Analysis Consortium (CPTAC) Biospecimen Core Resource and stored at −80°C prior to analysis.

### Sample processing

Cell line samples/tissues samples were lysed/homogenized in lysis buffer [8 M urea, 1x HALT Protease/Phosphatase Inhibitor Cocktail (Thermo Scientific)]. Lysates were cleared by centrifugation at 5000 g for 5 min at 4°C and protein concentration was measured by bicinchoninic acid assay (Pierce). Proteins were reduced with 10 mM dithiothreitol for 30 min at 56°C, alkylated with 55 mM iodoacetamide for 45 min at room temperature (RT) protected from light, and diluted 4-fold with 100 mM ammonium acetate, pH 8.9. Proteins were digested with sequencing grade modified trypsin (Promega) at an enzyme to substrate ratio of 1:50 overnight at RT. Enzymatic activity was quenched by acidifying with glacial acetic acid to 10% of the final solution volume, and peptides were desalted using C18 solid phase extraction cartridges (Sep-Pak Plus Short, Waters). Peptides were eluted with 60% acetonitrile in 0.1% acetic acid, dried using vacuum centrifugation, lyophilized in 800 μg aliquots, and stored at −80°C until analysis.

### Tyrosine phosphorylated peptide enrichment

Lyophilized tryptic peptide aliquots were resuspended in 400 μL of immunoprecipitation (IP) buffer [100 mM Tris-HCl, 0.3% NP-40, pH 7.4] and supplemented with a mixture of ∼1 pmol of each IS peptide standard. The light/heavy peptide mixture was incubated with 60 μL protein G agarose bead slurry (Calbiochem) conjugated to an antibody cocktail containing 12 μg 4G10 (Millipore), 12 μg PT66 (Sigma) and 6 μg of pY100 (Cell Signaling Technologies), rotating overnight at 4°C. Of note, samples 1 and 2 of the A549 enrichment analysis were only incubated for 6h at 4°C, whereas sample 3 followed the described protocol with overnight incubation.

Beads were washed 1x with IP buffer, 3x with 100 mM Tri-HCl, pH 7.4, and eluted in 2 rounds of 25 μL 0.2% TFA. Phosphopeptides were further enriched using High-Select Fe-NTA Phosphopeptide Enrichment Kit (Thermo Scientific) following manufacturer’s instructions with minor adjustments. Modifications include reducing the peptide volume initially added to the Fe-NTA column (50 μL) and reducing the elution volume to 2 rounds of 20 μL elutions. Peptide elutions were dried down using vacuum centrifugation to <2 μL total volume and resuspended in 5% acetonitrile in 0.1% formic acid for a total volume of 10 μL.

### Peptide Synthesis

Peptides were purchased from Thermo Scientific Custom Peptide synthesis service. All synthetic peptides used in this study were produced as a PEPotec Custom Peptide Libraries using FMOC solid-phase technology. The peptides were synthesized with the following specifications: crude purity, synthetic isotope-labeled c-terminal lysine (K) or arginine (R) or proline (P) or alanine (A) or isoleucine (I) or valine (V). The crude peptides after synthesis were dissolved in 0.1% TFA in 50% (v/v) acetonitrile/water and stored at −20 °C. A pool of first heavy peptide mixture was prepared by mixing an equimolar amount of each peptide with the final concentration at 1pmol/µl in 0.1% TFA and 3% (v/v) acetonitrile and subjected to nanoLC-MS/MS analysis to determine the intensity response of 340 peptides. A final heavy peptide mixture was prepared by increasing the concentration of the 58 “low-intensity” heavy peptides with low intensity response values. The final concentration of 58 heavy peptides was ranged from 1.8 to 5.5 pmol/µl. Exact concentrations are specified in data file S1.

### LC-MS/MS Analysis

Samples were analyzed using an Orbitrap Exploris 480 mass spectrometer (Thermo Scientific) coupled with an Easy-nLC 1200 (Thermo Scientific), Nanospray Flex ion source (Thermo Scientific), and column oven heater (Sonation). A 10 μL injection volume of sample was directly loaded onto a 25 cm Aurora Series emitter column (IonOpticks) with a column oven temperature of 40°C. Peptides were eluted at a flow rate of 400 nL/min across a linear gradient consisting of 0.1% formic acid (buffer A) and 80% acetonitrile in 0.1% formic acid (buffer B). The gradient is as follows: 3-19% B from 1-37 mins, 19-29% B from 37-51 mins, 29-41% B from 51-60 mins, 41-95% B from 60-63 mins, and 95-3% B from 70-70:05 mins.

### Survey MS analyses

A flowchart describing the pTyr SureQuant method build and analysis workflow can be found in fig. S5. Prior to SureQuant acquisition, the IS peptides were first characterized by data dependent acquisition (DDA) with an inclusion list of the precursor ions under +2, +3, and +4 charge states for each IS trigger peptide to select optimal charge states and product ions for subsequent targeted experiments. For this analysis, a mixture containing approximately 700 fmol of each IS peptide was directly injected. Next, a “survey run” was performed, still based on directed DDA but with an inclusion list focused on the optimal charge state for each peptide, to capture the precursor ion intensity responses and derive the intensity thresholds for MS^2^ scan triggering in subsequent SureQuant analyses. To take sample losses from pTyr enrichment steps into account in determining triggering thresholds, a nominal amount (∼1 pmol) of each IS peptide was added to 800 ug of the A549 processed cell line standard, followed by pTyr enrichment as a representative sample. Parameters obtained in these survey analyses were used in all subsequent pTyr SureQuant analyses.

The mass spectrometry parameters used for these preliminary analyses were as follows: spray voltage: 1.9kV, no sheath or auxiliary gas flow, heated capillary temperature: 280°C. DDA analyzes collected full-scan mass spectra with m/z range 300-1200, AGC target value: 1000%, maximum injection time (IT): 50 ms, resolution: 120,000. For every scan, the top 40 most intense ions on the inclusion list (if above a 1e5 intensity threshold) were isolated [isolation width of 1.0 m/z] and fragmented [normalized collision energy (nCE): 28%] by higher energy collisional dissociation (HCD), scan range: 100-1700 m/z, maximum IT: 10 ms, AGC target value: 1000%, resolution: 7,500.

### Targeted MS analyses for A549 and tumor samples

The SureQuant method combines various scan events and filters, depicted in fig. S6A. During SureQuant analyses, a high resolution MS1 scan is acquired to monitor the predefined optimal precursor ions of the IS heavy peptides, based on the list of associated m/z values and intensity thresholds. If any targeted m/z from the inclusion list is detected and meets the minimum intensity threshold specified, a short fill time, low resolution MS^2^ scan of the IS peptide is performed in the subsequent MS cycle. If the scan contains at least 5 of 6 specified product ions, a high resolution MS^2^ scan of the endogenous peptide at the defined mass offset is performed with longer fill times to improve measurement sensitivity. In the current implementation of SureQuant acquisition on the Exploris 480 MS, all trigger peptides scans are performed first, followed by target peptide scans in any given MS cycle in order to optimize parallelization of trapping and Orbitrap-FT processing in sequential scans (fig. S6B).

To implement this method, the custom SureQuant acquisition template available in Thermo Orbitrap Exploris Series 1.0 was utilized. The template is structured such that the acquisition parameters for each unique isotopically labeled amino acid and charge state (defining the m/z offset) is contained within a distinct 4-node branch stemming from the full scan node (fig. S6C). We utilized the default template, which contains 6 branches for the +2, +3, and +4 charge states of SIL lysine and arginine residues and added four additional branches for the +2 charge states of SIL proline, valine, isoleucine, and alanine for a total of 10 branches. In each branch, the peptide m/z and intensity thresholds are defined in the “Targeted Mass” filter node. Next, parameters for the low resolution, IS peptide MS^2^ scan are defined, followed by the “Targeted Mass Trigger” filter node, which defines the 6 product ions used for pseudo-spectral matching. To connect each set of product ions within the targeted mass trigger node to a given precursor mass, we utilize the group ID feature to define the precursor m/z associated with each group of product ions is related to. Finally, along with the scan parameters for the second MS^2^ scan of the endogenous peptide, we define the isolation offset (m/z) within each node.

Standard mass spectrometry parameters for SureQuant acquisition are as follows: spray voltage: 1.5kV, no sheath or auxiliary gas flow, heated capillary temperature: 280°C. Full-scan mass spectra were collected with a scan range: 300-1500 m/z, AGC target value: 300%, maximum IT: 50 ms, resolution: 120,000. Within a 5 second cycle time per MS1 scan, heavy peptides matching the m/z (within 10 ppm) and intensity threshold defined on the inclusion list were isolated [isolation width of 1.0 m/z] and fragmented [nCE: 28%] by HCD with a scan range: 100-1700 m/z, maximum IT: 10 ms, AGC target value: 1000%, resolution: 7,500. A product ion trigger filter next performs pseudo-spectral matching, only triggering an MS^2^ event of the endogenous, target peptide at the defined mass offset if n ≥ 5 product ions are detected from the defined list. If triggered, the subsequent light peptide MS^2^ scan has the same CE, scan range, and AGC target as the heavy trigger peptide, with a higher maximum injection time and resolution (for example, max IT: 180, resolution, 60,000), however these parameters vary slightly across samples in order to optimize acquisition speed and sensitivity.

### Internal standard peptide characterization and survey run analysis

All analyses were processed using Skyline software (*56*). The IS peptides properly detected in initial directed DDA analysis, *i.e.*, those for which at least one precursor ion yielded several MS^2^ scans including at least 6 theoretical y- or b-type fragment ions, were retained for SureQuant method development. For peptides detected under multiple charge states, only the precursor ion yielding the highest signal response was retained. For each peptide, 6 associated optimal fragments ions were selected for psudo-spectral matching (typically the most intense ones showing sufficient specificity, *i.e.*, without neutral loss or low m/z value). Individual intensity thresholds for each IS peptide was set to 1% of the precursor MS1 intensity value at the apex of its chromatographic profile in the survey run analysis.

### Mass spectrometry targeted pTyr analyses

Peak area ratios of endogenous light peptides and corresponding heavy IS peptides for the 6 selected product ions were exported from Skyline, and peptides were filtered according to the following criteria: First, only IS peptides with an AUC > 0 for n ≥ 5 product ions were considered. Of these remaining targets, only endogenous targets with an AUC > 0 for n ≥ 3 product ions were considered. For quantification, the peak area values of the 3 highest intensity product ions present for both the light/heavy peptides were summed, and the ratio of light endogenous to heavy IS peptide signal was taken across samples. We selected 3 product ions for quantitation to balance specificity with the ability to retain lowly abundant targets. For tumor sample analysis, L:H ratios of peptides quantifiable in all tumors were included in the full matrix, and peptides quantifiable in ≥ 16/31 tumors were included in the expanded matrix. Both matrixes were z-score normalized for specified analyses. Analyses were performed using Python 3.6.0.

### Protein expression profiling

LC-MS/MS raw files from the TMT-labeled global proteome analysis performed by Vasaikar et al. (*15*) were re-processed using Proteome Discoverer (version 2.2). All mass spectra were searched using Mascot (version 2.4) against the human SwissProt database with a tryptic enzymatic digestion, allowing for 2 missed cleavages, +/- 10 ppm parent ion tolerance. Static modifications of Cys carbamidomethylation and TMT on N-terminus and Lys residues were included, along with variable Met oxidation. Peptide spectrum matches (PSMs) were filtered according to the following criteria: Search engine rank = 1, isolation interference ≤ 30, ion score ≥ 20, peptide length ≥ 6. Relative protein abundance was calculated as the ratio of tumor abundance to reference channel abundance (TMT-131) using the summed TMT reporter ion intensities from all peptides uniquely mapped to a gene. Relative abundances were next divided by the median relative abundance ratio from each TMT channel to correct for sample loading variation within each analysis. Adjusted relative abundances for proteins quantified across all 31 tumors were z-score normalized for subsequent analyses.

### RNA-sequencing

RNA-sequencing data was analyzed by Vasaikar et al., as previous described (*15*). RSEM upper-quartile normalized values for the tumor panel used in this study were extracted and z-score normalized for subsequent analyses.

### Principal component analysis

PCA was performed in Matlab R2019b using z-score normalized L:H ratios of the 26 tumors with a defined unified molecular subtype. Only peptides identified across all tumors were used (165 unique sites).

### Enrichment analysis

For tumor-specific pathway enrichment analyses (TPEA), Source proteins of phosphorylated peptides were rank ordered from highest to lowest z-score. In cases where more than one peptide mapped to the same source protein, the maximum/minimum was selected, depending on the directionality of the enrichment analysis. We utilized gene set enrichment analysis (GSEA) 4.0.3 (*57*) pre-ranked tool against a custom database of 12 pathways (data file S3), obtained from gene ontology (GO) biological processes terms, Reactome pathways, and KEGG pathways with 1000 permutations, weighted enrichment statistic (p=1), and a minimum gene size of 12. Results were filtered according to p < 0.05, FDR q-value < 0.25.

Similarly, for kinase-substrate enrichment analysis (KSEA) all peptides were ranked ordered by z-score and pre-ranked GSEA was performed using a custom library of 12 phosphosite specific kinase-substrate sets (data file S3) from the Substrate Kinase Activity Inference (SKAI) library (*30*) using parameters listed above and a minimum gene set size of 10. Results were filtered according to p < 0.1 and datasets with FDR q-value < 0.25.

GSEA using RNA-sequencing and protein-expression profiling data was similarly performed by rank-ordering genes by z-score and analyzed against the Molecular Signatures Database hallmarks gene sets with parameters listed for TPEA and a minimum gene size of 15. Results were filtered according to p < 0.05, FDR q-value < 0.25.

### Protein-protein interaction network analysis

Significantly enriched pathways and biological processes (FDR q-value < 0.05) were identified within clusters of co-regulated phosphopeptides using STRING v11 and visualized using Cytoscape v3.7 (*58, 59*). Nodes are annotated by pTyr peptide gene name, and edges represent protein-protein associations experimentally determined.

## Supplementary Materials

Fig. S1. pTyr sites measured in human colorectal tumors.

Fig. S2. Correlation analysis identifies tumors with significant pathway enrichment.

Fig. S3. ErbB signaling pathway enrichment analysis.

Fig. S4. Tumor-specific pTyr signatures of T cell signaling peptides.

Fig. S5. Flowchart of SureQuant pTyr workflow.

Fig. S6. SureQuant pTyr acquisition method framework and parameters. Data file S1: Targeted tyrosine phosphorylated peptides.

Data file S2: Clinical data for tumor specimens analyzed.

Data file S3: Custom pathway and substrate-kinase libraries for tumor-specific pathway enrichment analyses.

## Acknowledgements

We thank Andreas Huhmer at Thermo Scientific for funding the synthesis of the stable isotope labeled peptide standards, and for providing project guidance and manuscript feedback; The NCI Clinical Proteomic Tumor Analysis Consortium (CPTAC) for providing tumor samples and for data analysis guidance with RNA-sequencing and proteomics data (Bing Zang); Brian Joughin for data analysis recommendations.

## Author contributions

Project concept and design: LES, ASG, BP, SG, FMW. Methodology development: LES, CTF, ASG, BP, SG. Data acquisition: LES, CTF, ASG, BP. Analysis and interpretation of data: LES, CTF, ASG, BP, SG, FMW. Writing, review, and revision of the manuscript: LES, CTF, ASG, BP, SG, DL-F, FMW. Project advising: DL-F, FMW.

## Supplementary Materials

**Fig. S1.**
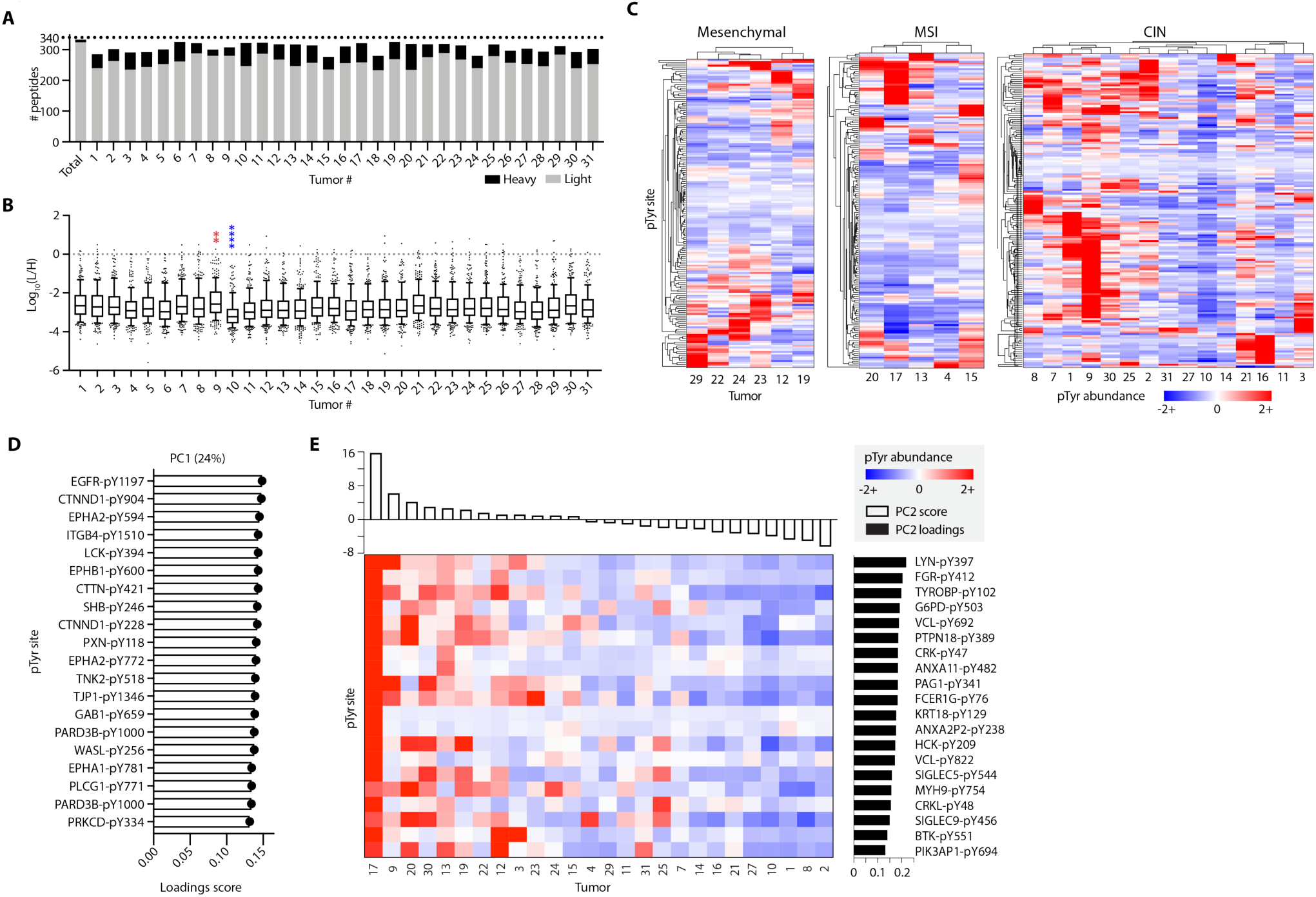
pTyr sites measured in human colorectal tumors. (**A**) Number of unique heavy (black) and light (grey) peptides identified in each tumor. (**B**) Distributions of light to heavy signal ratios (L/H) for each tumor. Data is displayed as a box and whiskers plot, where the box describes the interquartile range and the whiskers define the 10-90 percentile of data. * Indicates significantly increased (red) or decrease (blue) from the mean distribution using Dunnett’s multiple comparison test for significance. **p<0.01, ****=p<0.0001. (**C**) Hierarchical clustering of pTyr peptides within each unified multi-omics subtype defined by Vasaikar et al. Distance metrics for clustering of peptides and tumors, respectively, are Euclidean and correlation. (**D**) Peptides with the top 20 loadings scores for PC1, pY denotes the residue position with pTyr modification. (**E**) Top 20 loadings scores of the peptides derived from unique proteins for PC2, ranked from highest to lowest PC2 score with corresponding pTyr abundance levels (z-score normalized L/H) across tumors.

**Fig. S2.**
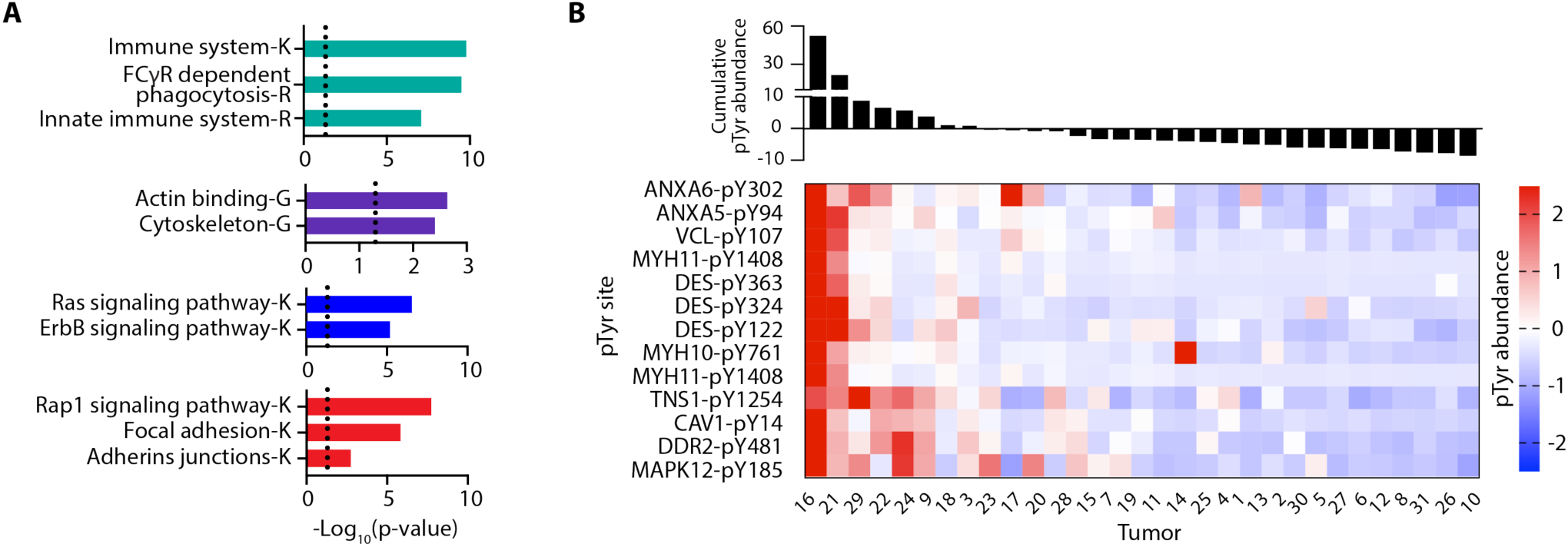
Correlation analysis identifies tumors with significant pathway enrichment. (A) Selected significantly enriched pathways identified in Fig. 4. K=Kegg pathway, G=Gene Ontology term, R=reactome pathway. Significance values are FDR-adjusted, and a cutoff of p < 0.05 was used for filtering. (B) pTyr peptides from cluster 2, rank ordered from highest to lowest cumulative pTyr abundance, where abundance values are z-score normalized light to heavy signal ratios.

**Fig. S3.**
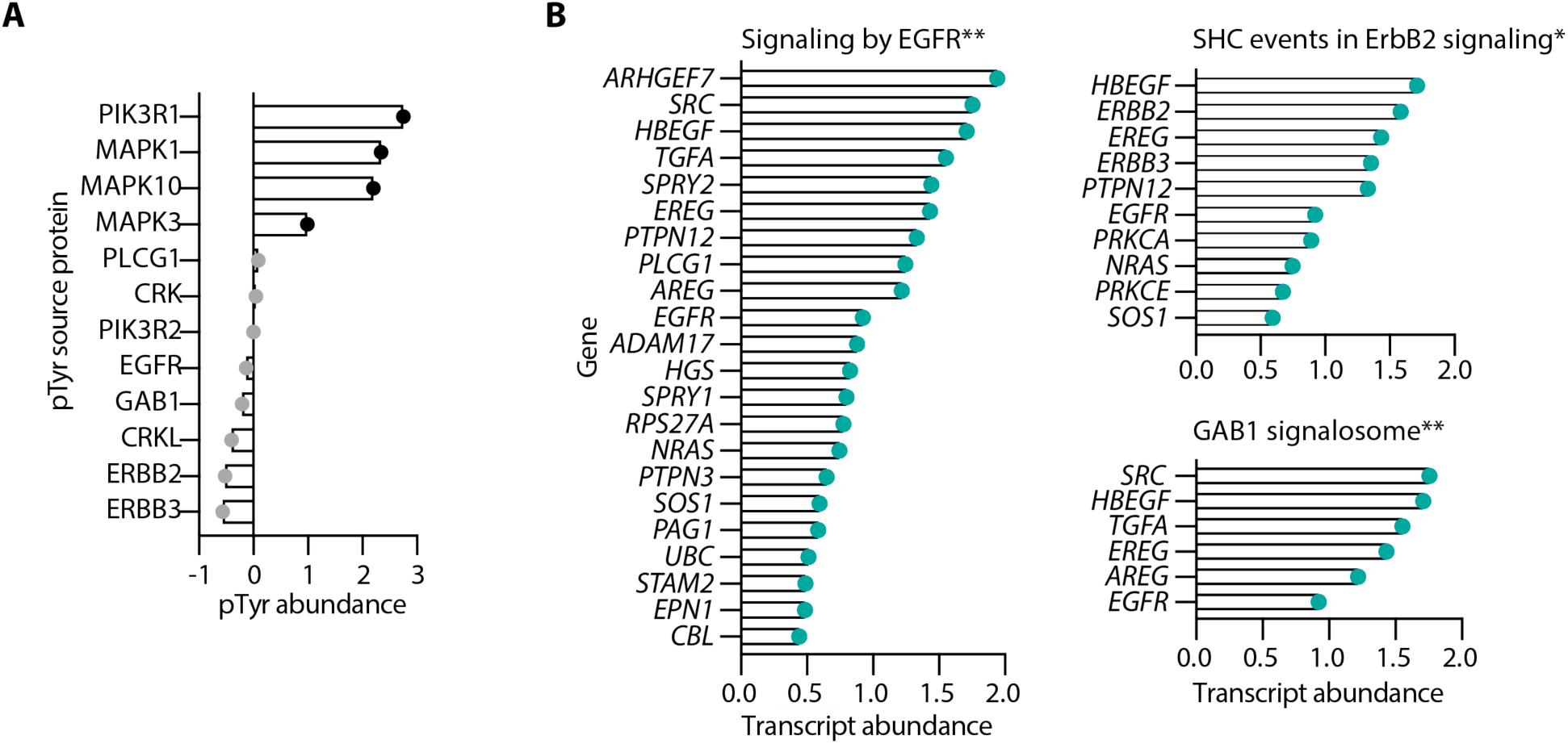
ErbB signaling pathway enrichment analysis. (A) Tumor 19 pTyr abundance of peptides in the ErbB signaling pathway. All peptides are annotated by source protein, with peptides in the enrichment core colored in black. (B) Tumor 9 positively enriched reactome pathways with corresponding transcript abundance levels (z-score normalized). *= p<0.05. **=p<0.01 FDR q-value < 0.05 for all.

**Fig. S4.**
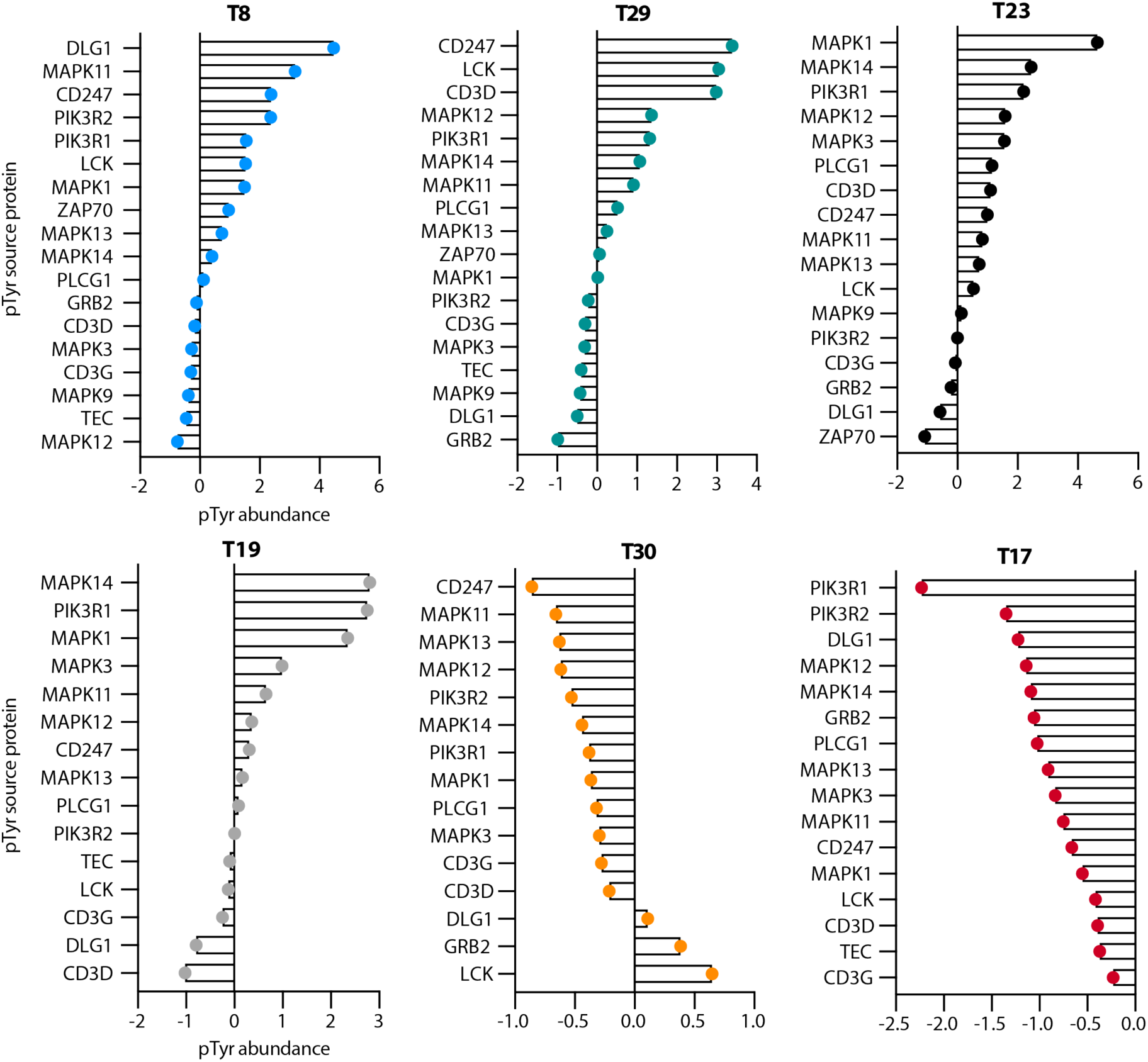
Tumor-specific pTyr signatures of T cell signaling peptides. pTyr abundance (z-score normalized light to heavy signal ratios) of T cell signaling peptides in tumors with significant pathway enrichment. Peptides are annotated by source protein.

**Fig. S5.**
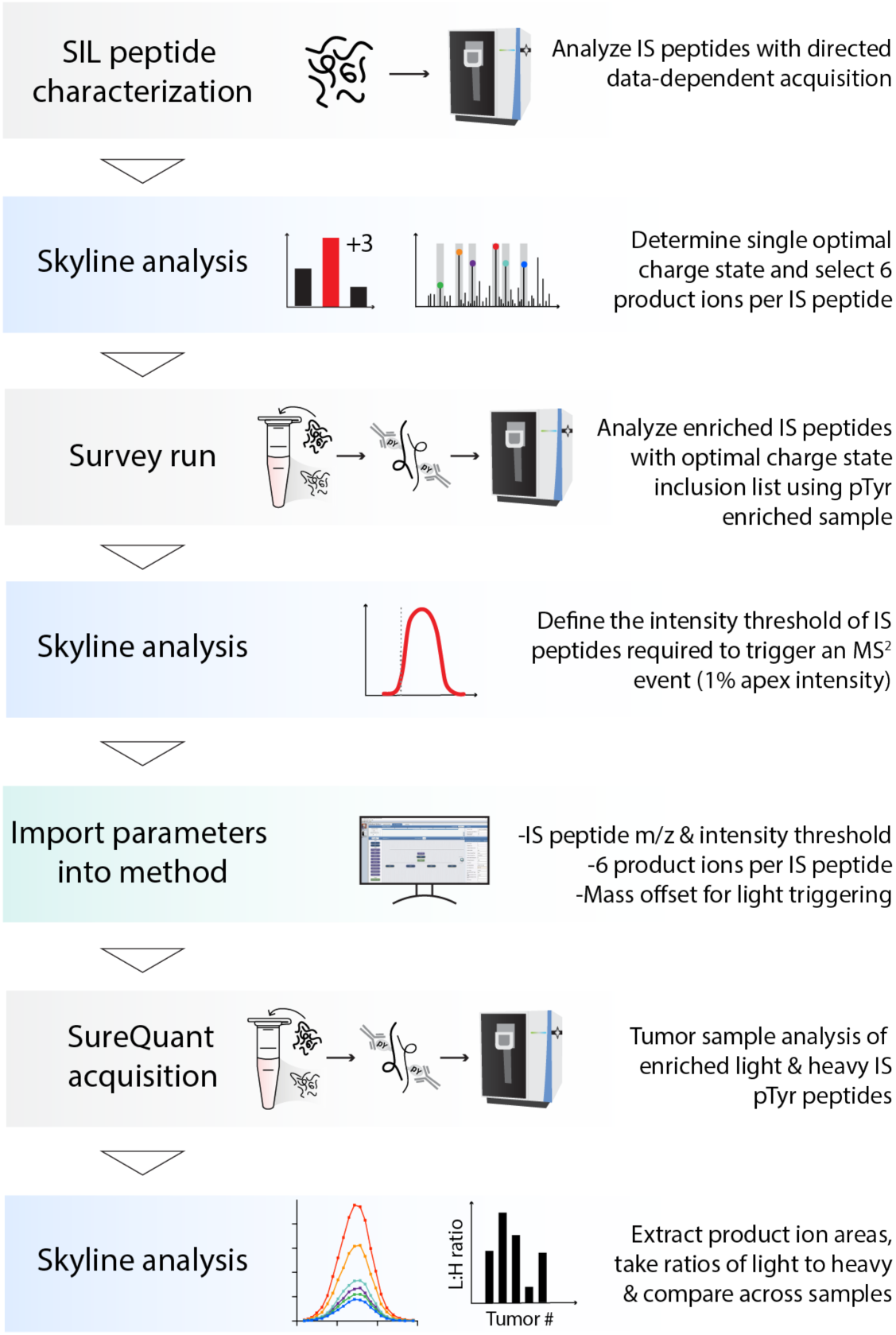
Flowchart of SureQuant pTyr workflow. SureQuant pTyr pipeline for method building, data acquisition, and data analysis.

**Fig. S6.**
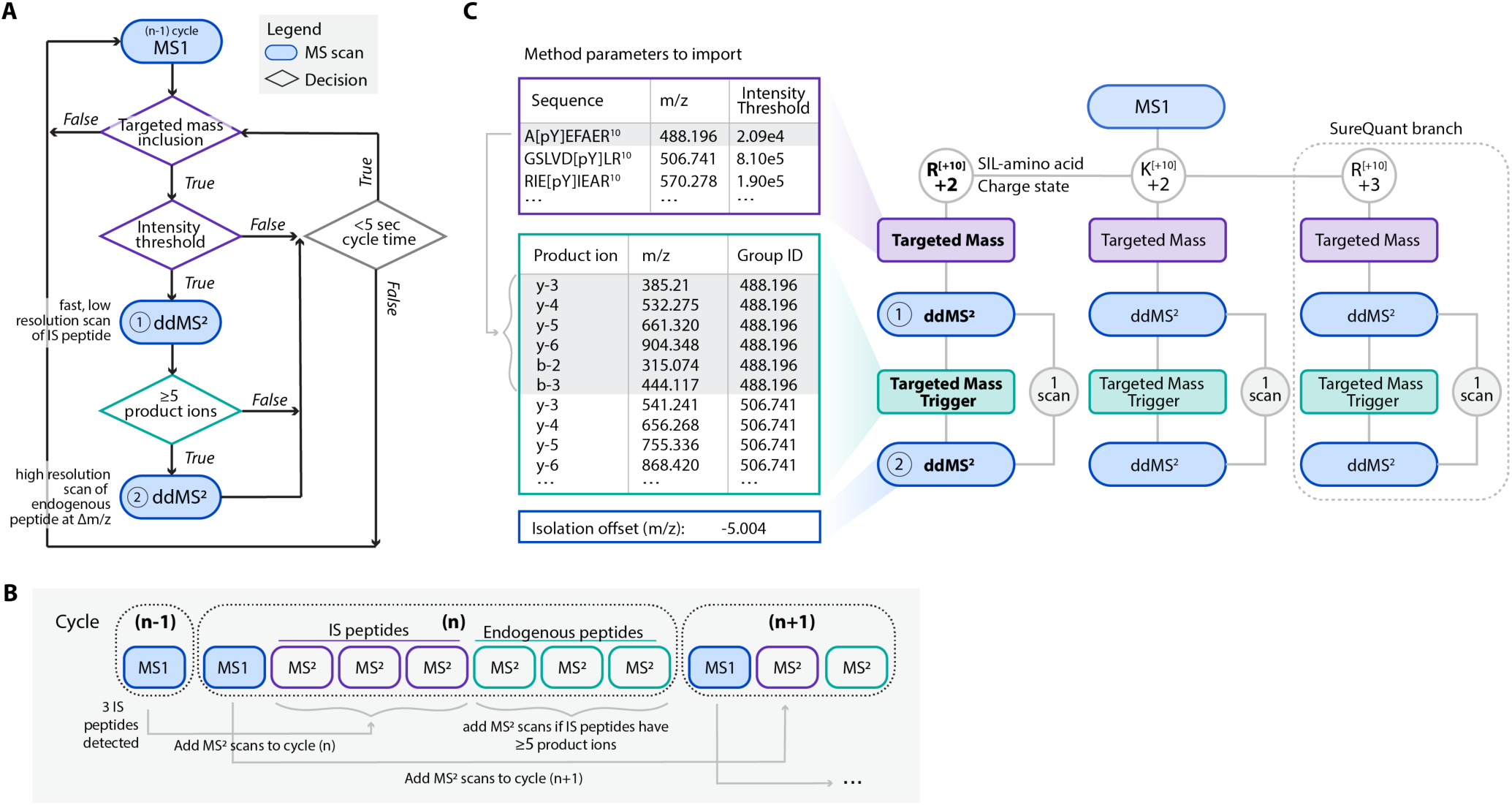
SureQuant pTyr acquisition method framework and parameters. (**A**) Decision tree for SureQuant acquisition method framework. (**B**) Scan sequence for SureQuant analyses on the Exploris 480 MS. For example, if three internal standard (IS) peptides are detected in the MS1 scan of cycle (n-1), an MS^2^ scan of each IS peptide is added to the scan sequence in cycle (n). If each IS MS^2^ scan contains ≥5 pre-defined product ions, MS^2^ scans of each endogenous peptide at the defined m/z offset are added to the scan sequence in cycle (n). This scan structure repeats for each cycle. (**C**) Method scan structure and parameters to import for SureQuant acquisition illustrated for three branches, however SureQuant pTyr uses ten branches. [pY] denotes the residue position with pTyr modification, and R^10^ and K^10^ denote stable isotope labeled (SIL) arginine (+10) and lysine (+10), respectively.

